# Tracing evolution history of 100 whole genome sequences of diffuse stem brain tumor

**DOI:** 10.1101/2021.03.14.435347

**Authors:** Lihua Zou

## Abstract

Diffuse intrinsic pontine glioma (DIPG) is a deadly disease among young children. The evolution path and mutational processes giving rise to DIPG remain elusive. We analyzed 100 whole genome sequences (WGS) from 60 DIPG patients. This revealed 25% DIPGs acquired whole-genome duplications (WGD) early during tumor evolution. WGD samples are associated with loss of TP53 and poorer survival. In addition, almost all WGD samplers harbor complex structural variations (SVs) and show characteristic short microhomology at SV breakpoints. Mutation analysis revealed that H3K27M driver mutation is acquired early during tumor clonal evolution. Mutation signature analysis identified a unique mutational process at a late stage of tumor evolution. This study revealed that tumor evolution of DIPG is characterized by chromosomal instability shaped by DNA repair defects and dynamic mutational processes. Our work shed new insights on the disease pathogenesis of DIPG and provided rationale for designing novel therapy for this deadly disease.

## Introduction

Diffuse intrinsic pontine glioma (DIPG) is an incurable pediatric brain tumor with a median survival less than twelve months[1]. High-throughput sequencing has shown nearly 80% of DIPGs carrying a specific mutation resulting in replacement of lysine 27 by methionine(K27M) in the encoded histone H3 proteins[2,3]. Large-scale sequencing analysis further defined the molecular diversity among DIPGs and identified co-segregating mutations within histone-mutant subgroups[4,5]. It is emerging that DIPG is a much more heterogeneous disease than previously thought.

Although sequencing analysis have identified drivers in DIPGs, little is known about the evolution dynamics of DIPG. WGD, involving the doubling of the whole chromosome complements, is a singular catastrophic event during tumor evolution and thought to have a profound impact on tumor evolution. A recent pan-cancer analysis has shown WGD is a common genomic hallmark of human cancer[6,7]. The frequency of WGD is highly variable and present most commonly in cancer types with high mutation rates such as lung squamous cell carcinoma and triple-negative breast cancer[7]. Previous work was based on mostly adult cancer, the extent of WGD in childhood cancer, however, has not been systematically characterized.

To characterize WGD and its impact on tumor evolution in DIPGs, we performed a meta-analysis to combine whole genome sequencing (WGS) data from 60 DIPG patients (see Methods). We performed copy number analysis to identify WGD samples among DIPG samples. We examined genomic features that correlated with WGD. We conducted computational analysis to infer the timing of WGD, H3K27M and other driver events during tumor evolution. We performed mutation signature analysis to characterize dynamic mutation signatures during tumor evolution.

## Results

### Genome duplication is a genomic hallmark of DIPG

To quantify the frequency of genome doubling in DIPG, we combined whole genome sequencing data based on 60 patients enrolled from two independent cohorts (see Methods). We applied an established statistical measure to quantify genome duplication based on fraction of allelic gain on autosomal chromosome[6]. This revealed that 25% of patients of our study underwent extensive genomic gain with allelic gain present on over 50% of their autosomal tumor genome (Figure 1B), which is consistent with the genomic pattern of WGD reported[6].

**Figure 1:**
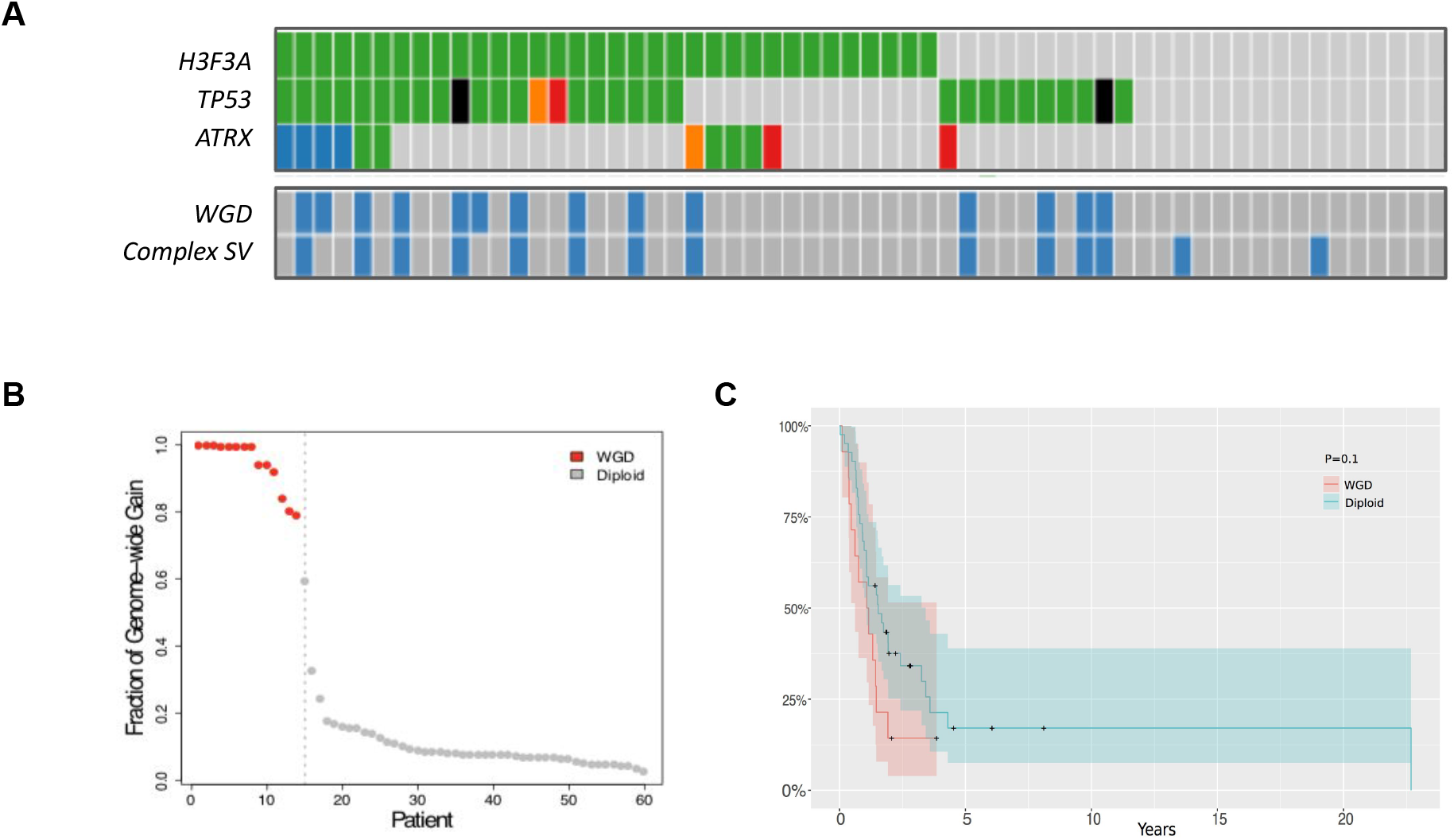
Whole genome duplication is a genomic hallmark of DIPG. A) DIPG driver mutations vs WGD and complex SV status sample-by-sample. B) Sample classification based on fraction of genome-wide gain as indicated on y-axis. C) Kaplan-meier survival analysis indicates WGD samples have significantly worse outcomes than other patients in the study.

To correlate WGD with molecular features, we compared the association between WGD and mutation status of a list of common DIPG drivers. This indicated TP53 mutation is enriched among WGD patients (Figure 1A; pval < 1e-6; Chi-Square test). This supports the experimental evidence TP53 is required to prevent genome-doubled cells from re-entering the cell cycle and proliferation[8,9]. We found nearly all WGD patients have shown patterns of complex structural variations (SV), which include 11 patients shown pattern of chromothripsis and 3 patients who have high copy numbers changes linked across multiple distant chromosome segments which suggest ecDNA (e.g. double minute, neochromosome) as underlying structure (Figure 1A, 2A). In addition, Kaplan-meier survival analysis indicated WGD patients have significantly worse outcomes than non-WGD patients (Figure 1C; p<0.1).

**Figure 2:**
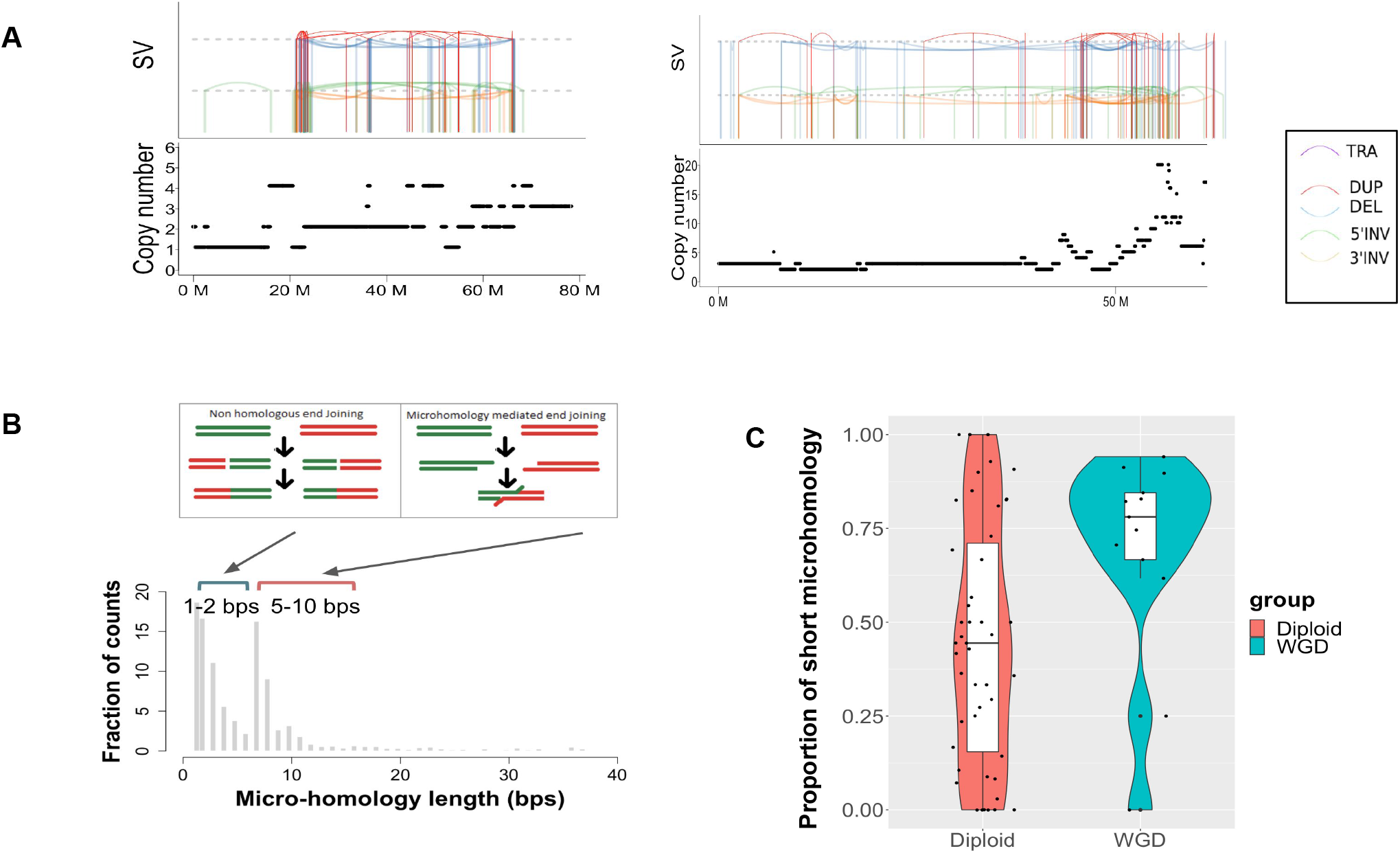
Genome duplication is associated with error-prone DNA repair mechanisms. A) Examples of complex SVs including a patient (left panel) carrying chromothripsis and another patient (right panel) carrying ecDNA in their genome. Colored lines in the upper part of each panel indicate different types of SVs (see legend). Line segments in the bottom part of the panel indicate the total copy number along the chromosome. B) Bimodal distribution of micro-homology length at junctions of SV breakpoints; x-axis shows bins of micro-homology length; y-axis shows fractions of breakpoint counts within each bin. C) Distribution of short micro-homology associated breakpoints in WGD (green) and Diploid (red) samples. y-axis in the boxplot shows the proportion of short micro-homology associated breakpoints in each sample.

### Genome duplication is associated with error-prone DNA repair mechanisms

We found almost all WGD patients (13/14) harbored complex SVs (Figure 1A). Figure 2A showed two examples of complex SVs identified in our study. SV is the result of mis-repaired DNA. To gain insight about the mutational process underlying SVs in DIPG, we studied the DNA micro-homology sequence at breakpoint junctions of SVs. This revealed a bimodal distribution based on micro-homology at SV breakends (Figure 2B) including a peak with short micro-homology length centered on 1-2bps and a second peak with long micro-homology length between 5-10 bps. We further performed an analysis to quantify the proportion of short vs long micro-homology in each sample from our study. This unraveled a significant enrichment of short micro-homology surrounding SV breakpoints among WGD patients (Figure 2C). Previous studies showed that short 1-2bps micro-homology at breakpoints are typically associated with error-prone Non-homologous End-Joining DNA repair processes (NHEJ)[10]. Taken together, our analysis suggests error-prone DNA repair as a mechanism underlying excessive mis-repaired SVs in WGD DIPG patients.

### Dynamic mutational processes underlies ongoing tumor evolution

Mathematical methods modeling genome-wide mutations have linked specific DNA repair defects with deficiencies in the homologous recombination, mismatch repair and nucleotide-excision repair (NER) pathways[11–15]. We performed mutation signature analysis and identified 4 distinct mutation signatures from our study. This include a signature characterized by C>T transitions at CpG dinucleotide due to age-related accumulation of 5-methylcytosine deamination events (Sig.1), a signature exhibiting a mutation pattern associated with alkylating agents possibly due to treatment (Sig.11), and a signature closely resembles Sig.5 which is present in all cancer types and that has transcriptional strand bias for T>C substitutions[16]. Interestingly, there is a strong statistical association between Sig.5 and H3K27M (p<1e-4) (see Supplementary materials). This association is not seen for other DIPG drivers including ACVR1 and TP53 suggesting unique mutation processes operative in H3K27M DIPG.

In addition, we identified an unknown signature (Sig.U) that shares some similarity to Sig.3 and Sig.8 as previously reported in the COSMIC mutation signature database (cosine similarity 0.5-0.6) (Figure 4A)[16]. To further investigate the origin of this signature, we separated the mutations into clonal and subclonal mutations and repeated the analysis. This showed that the unknown mutation signature consists mostly of subclonal mutations (Figure 3B,C). This suggests the mutation spectrum is dynamic and can shift due to ongoing tumor evolution.

**Figure 3:**
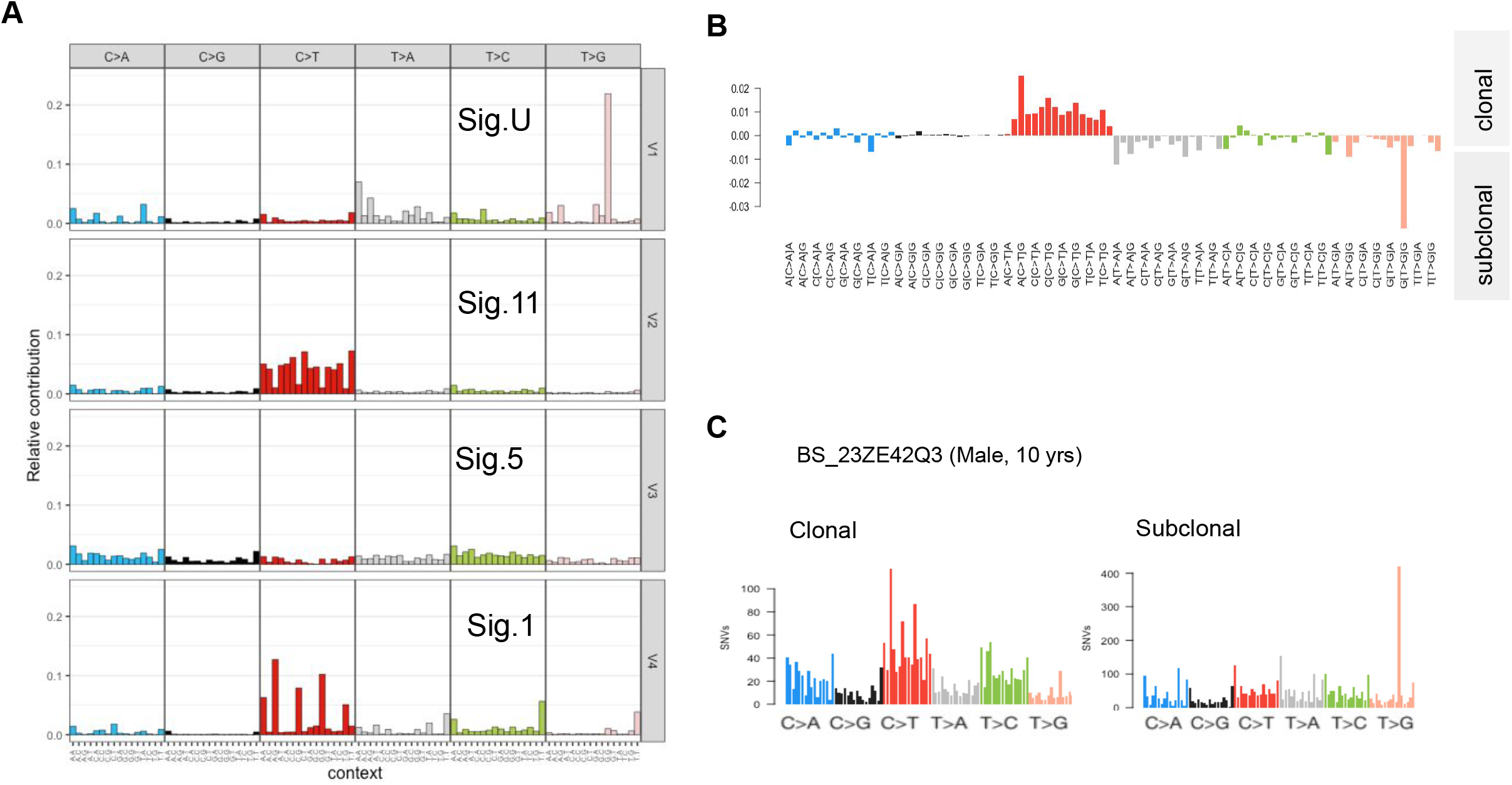
Mutation signatures of DIPG. A) Each panel shows a mutation signature identified in our study. This includes an unknown signature (Sig.U) in the top panel and Sig.1, Sig.5, Sig.11 as previously reported. x-axis shows tri-nucleotide context; y-axis shows relative contribution of each mutation context. B) Each bar shows the net proportion of mutations (clonal vs subclonal) at each of the trinucleotide contexts. Net proportion (indicated on y-axis) is calculated by subtracting the number of subclonal mutations from clonal mutations at each trinucleotide context. C) An example showing a patient has a different mutation spectrum between clonal and subclonal mutations.

**Figure 4:**
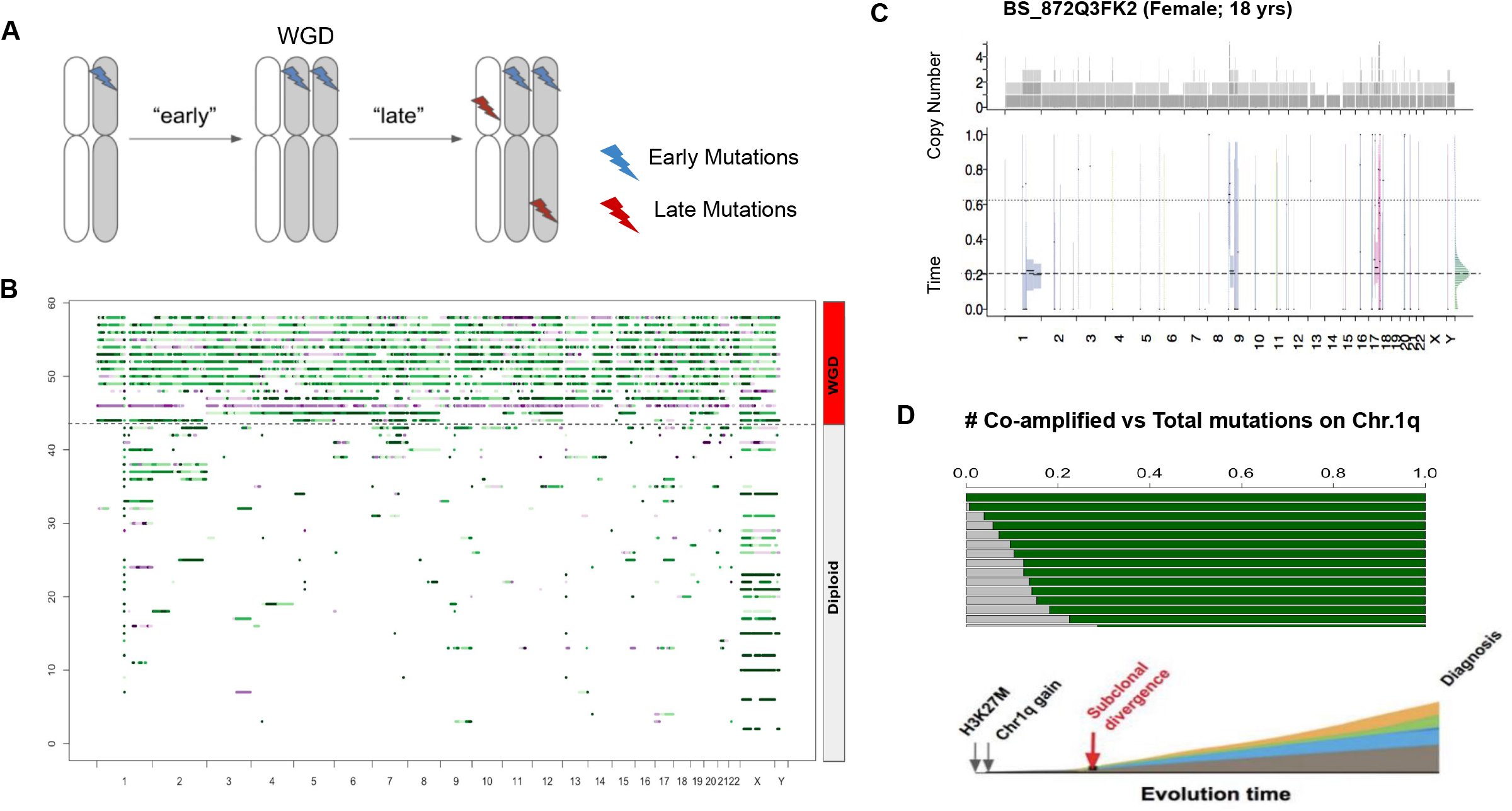
Timing of landmark events of DIPG. A) An illustration of “early” vs “late” mutations relative to WGD. Early mutations got co-amplified due to WGD while late mutations are present in a single copy. B) Timing of chromosomal gain in each patient. Each line segment indicates a chromosomal gain and the color indicates the chromosomal gains happened ‘early’ (green) or “late” (purple). C) An example showing a patient had copy number gain of Chr.1q early during tumor clonal evolution. Top panel shows allele specific and total copy numbers across the chromosomes; the y-axis of the bottom panel shows relative mutational time during clonal evolution. D) Timing of H3K27M. Each bar in the top panel indicates a sample with both H3K27M and Chr.1q gain; the length of grey bar indicates proportion of co-amplified mutations (out of all clonal mutations) in a sample. Bottom panel illustrates a projected timeline of landmark events in a patient with H3K27M and Chr.1q gain.

### Timing of WGD and K27M mutation

Somatic mutations accumulated in individual cancer genomes provide a record of the mutation history of tumor evolution[7,17,18]. Mutations co-amplified on the duplicated DNA segments must have occurred before the copy number gained[19]. This temporal relationship can be inferred from the mutant allele frequency, after adjusting for tumor purity and ploidy to obtain the absolute number of alleles carrying the mutation[19–21]. We sought to investigate the relative timing of WGD during the tumor evolution of DIPG. We followed the computational framework[7] using WGS data, where the proportion of mutations co-amplified on a chromosomal gain region is used as a proxy to distinguish “early” vs “late” events (Figure 4A). Our analysis showed overall there are relatively few mutations co-amplified within the chromosomal gain region. This suggests that WGD and most copy number gains identified in our study are acquired early during tumor evolution (Figure 4B).

Next we sought to estimate the timing of H3K27M mutations from our study. H3K27M was present in nearly 80% DIPG patients and was thought to be the tumor initiation event in DIPGs[22]. H3K27M is present on Chr.1q. Our analysis showed Chr.1q is among the most frequent amplified events in our study (Figure 4B) and typically gained early during tumor evolution (Figure 4B,C). Furthermore, focusing on a subset of patients with both H3K27M and Chr.1q gain, our analysis showed that there are on average 15% co-amplified mutations along with H3K27M present on Chr.1q (Figure 4D; upper panel). This indicates H3K27M must be acquired before copy number gain of Chr.1q in those patients and before the majority of mutations accumulated during tumor clonal evolution (Figure 4D).

## Discussion

Taken together, we showed WGD is more prevalent than previously thought in DIPG. We showed TP53 is associated with WGD in DIPG. Our analysis showed there is a strong association between WGD and complex SVs. The large number of DNA breakpoints associated with complex SVs indicated chromosomal instability in these patients. Despite the fact that these patients are of young age, their WGD frequency is similar to adult brain tumor[6]. Our analysis indicated error-prone DNA repair mechanisms such as NHEJ may contribute to the larger number of DNA breakpoints with short micro-homology at breakpoint junctions. However, the detailed molecular machinery contributing to the chromosomal instability in DIPG is yet to be elucidated. We expect our study will pave the way for further studies to identify specific vulnerabilities for treating WGD DIPG patients.

Furthermore, our analysis showed that WGD and H3K27M are acquired early during tumor evolution. We showed that stage-specific mutational processes exist and the mutational processes can differ between early vs late stage of tumor evolution. This suggests the cooperation between a small set of drivers (e.g. H3K27M, TP53) and WGD can play a role in directing tumor evolution. Our study underscores the importance to take into consideration ongoing tumor evolution and tumor heterogeneity when treating this deadly disease.

## Methods

### Cohort description

The DIPG specimens used in our study are composed of radiologically diagnosed DIPG from Children’s Brain Tumor Tissue Consortium (CBTTC) and the Pediatric Pacific Neuro-oncology Consortium (PNOC). The raw whole genome sequencing and RNA-seq data can be downloaded from the Gabriella Miller Kids First Data Resource Center (KF-DRC)(https://kidsfirstdrc.org/).

### Variant calling of whole-genome sequences

The processed data can be downloaded from the CAVATICA(https://cavatica.squarespace.com/). Briefly, Somatic variants were processed using pipelines setup in the common workflow language described before [23]. Paired-end DNA-Seq reads were aligned to hg38 (patch release 12) reference genome using BWA-MEM. BAMs were merged and processed using Broad’s Genome Analysis Toolkit (GATK). Strelka2 was used to call Indels and Mutect2 was used to call SNVs using default parameters. The final Strelka2 and Mutect2 VCFs were filtered for PASS variants for downstream analysis. Manta was used to call structural variants (SV).

### Allelic DNA copy-number analysis

We used Sequenza[24] to generate allele-specific copy number calls from WGS data. In addition, Sequenza is applied to obtain the purity and ploidy estimates as well as integer copy-number calls at a given chromosomal segment for each sample. To classify samples based on WGD, we used a methodology from previous work[6]. A tumor is considered to have undergone WGD if there are over 50% of its autosomal genome have at least two copies of major alleles gained on a chromosomal segment.

### Mutation timing analysis

We infer the expected allele copy number of a mutation (MCN) as outlined previously[7]. Briey, MCN is a function of tumor purity (ρ), variant allele fraction (VAF), and total copy number (TCN). The formula is given by:

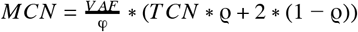

We classified mutations as outlined before[7]. Specifically, mutations present on >=2 copies/cell are classified as “early” clonal; mutations present on either on 1 copy/cell either on amplified or retained allele are classified as clonal; mutations present on <1 copy/cell are classified as subclonal. In the case of copy neutral LOH when no retained allele present, mutations present on 1 copy/cell is classified as “late” clonal. The proportion of early and late point mutations is used to estimate the relative timing of gained segments as outlined before[7].

### Mutation signature analysis

We extract the mutation signatures using MutationPattern[25]. Briefly, all mutations on the autosomal chromosomes were classified into 96 possible mutation types based on six base substitutions (C>A,C>G,C>T,T>A,T>C,T>G) within the tri-nucleotide sequence context including the immediate 5’ and 3’ base next to the mutated base. NMF algorithm is applied to decompose the mutation count into a set of representative signatures. We delineate the resulting sample-specific signature with a set of established mutation signatures documented in the COSMIC database using deconstructSigs[26].

### Structural variation analysis

We used ShatterSeek[27] to identify samples with inter- or intra-chromosomal clustering of SVs. We classified samples into types of complex SVs as outlined before[27,28]. We manually review samples to check the sequencing artifact, read coverage, and discordant and split read support at the discovered breakpoints.

## Acknowledgement

We thank members of Children’s Brain Tumor Tissue Consortium (CBTTC) (www.cbttc.org) for their support of open access, biospecimen driven research.

